# Negative Synergistic Effects of Drought and Heat During Flowering and Seed Setting in Soybean

**DOI:** 10.1101/2024.01.10.575108

**Authors:** Sadikshya Poudel, Ranadheer Reddy Vennam, Lekshmy V. Sankarapillai, Jinbo Liu, K. Raja Reddy, Nuwan K. Wijewardane, M. Shahid Mukhtar, Raju Bheemanahalli

**Affiliations:** Department of Plant and Soil Sciences, Mississippi State University, Mississippi State, MS, USA; Department of Biology, University of Alabama-Birmingham, Birmingham, AL, USA; Department of Agricultural & Biological Engineering, Mississippi State University, Mississippi State, MS, USA

**Keywords:** Interactive heat and drought, physiology, vegetative indices, seed quality, and gene expression

## Abstract

Rising temperatures and intense heatwaves combined with lower precipitations are the new norms of current global scenarios. These altered climatic conditions negatively impact soybean yield potential and quality. Ten soybean cultivars were subjected to four different growing conditions: control, drought, heat, and combined heat and drought to understand the physiological, yield, and molecular changes. Stomatal conductance was reduced by 62% and 10% under drought and heat, respectively. This reduction was further exacerbated to 93% when exposed to both stresses simultaneously. The highest canopy temperature was recorded at +8 °C with combinatorial treatments, whereas heat and drought exhibited +5.4 °C and +2 °C, respectively. Furthermore, combined stress displayed a more pronounced negative impact on greenness-associated vegetative index; the gene expression analysis further corroborated these findings. Particularly, each °C increase in temperature during flowering-seed filling reduced seed weight by ∼7% and ∼4% with and without drought, respectively. The seed protein increased under drought, whereas the oil showed a converse trend under drought and combined stresses. Most physiology and yield traits showed no significant correlations between control or individual and combined stress. This suggests that selection for combinatorial stress may not be appropriate based on nonstress or individual stress performance. Thus, incorporating stress-resilient traits into elite soybean cultivars could significantly boost soybean production under hot and dry climatic conditions.

## 1 INTRODUCTION

Soybean (*Glycine max* [L.] Merr.) is a leading oilseed crop grown in a wide range of climatic conditions (Li et al., 2017; Zhu et al., 2020). After corn, soybean is the most widely planted crop in the United States, accounting for 32% of the total cultivated land (2021-Soystats, 2021). Although the USA is the second largest soybean producer, >90% of the US soybean is produced under rainfed conditions. About 51% of soybean-growing regions in the US are exposed to drought (USDA Drought Monitor, 2023), and 80% of these are affected by heat stress (USDA ERS, 2023). Approximately 75% of the soybean acreage in Mississippi is rainfed (Zhang et al., 2016), with no supplement irrigation. Recent years have seen increased temperatures and extended periods of insufficient precipitation, contributing to a projected yield decline of up to 92% by 2050 (Yu et al., 2021b). Most southern US soybean-growing areas are exposed to these climatic conditions during critical growth stages. With an increased frequency of heatwaves and prolonged drought spells during the reproductive-seed fill stages, soybean production is predicted to face significant yield and quality losses. On the other hand, many studies have shown that the impact of combined stressors during the reproductive stages was greater than in the early vegetative stage in crops (Jumrani and Bhatia, 2018).

Exposure of soybean cultivars to drought stress during the reproductive and seed filling decreased seed number and weight (Poudel et al., 2023b). These decreases were associated with reduced stomatal conductance and increased canopy temperature under drought. Under heat stress, stomatal conductance and transpiration were increased or on par with control, but seed number (4.2%) and seed weight (5%) reduced per ℃ increase in temperature over 32 ℃ (Poudel et al., 2023a). For every one °C increase above average temperature (27.6°C), soybean yields are expected to decline by 2.4% (Hatfield et al., 2011; Alsajri et al., 2022). The increase in canopy temperature reduced CO_2_ assimilation by decreasing the leaf size and cell membrane stability (Onat et al., 2017). Under these stresses, chloroplasts experience oxidative damage (Vennam et al., 2023b). This primarily affects the photosynthetic process and accelerates leaf senescence by activating metabolic changes at the source (leaf) and sink (seed) (Hatfield et al., 2011; Wang et al., 2008). However, soybean genotypes respond differently to individual heat stress or drought, with some genotypes exhibiting significant plasticity (Poudel et al., 2023a). Small-seeded soybeans were less sensitive to heat stress than large-seeded soybeans (Puteh et al., 2013). Moreover, little is known about soybean’s genetic variability to combined stressors during the reproductive stage.

The intricate interplay of combined stressors threatens agriculture production. These interactive stressors elicit unique physiological, yield, and metabolic responses in plants, distinguishing them from responses triggered by individual stressors (Mittler, 2006; Prasch and Sonnewald, 2013; Cohen et al., 2021b). Previous studies demonstrated that the combined stress impact is much more significant and complex depending on crops’ growth stages (Schauberger et al., 2017; Matiu et al., 2017). Knowledge from other crops has shown the impact of hot or dry events on growth, reproduction, and grain-filling processes (Bheemanahalli et al., 2022b). Stressors that disrupt the crucial developmental stages of soybean, such as pollination, fertilization, and seed formation, cannot be mitigated in later growth stages (Krishnan et al., 2020; Poudel et al., 2023b; a). Notably, the interaction of drought and heat around flowering disrupts reproductive success and physiological-biochemical functions associated with seed filling. In the case of model plants such as Arabidopsis, exposure to combined drought and heat stress has been shown to cause a decline in sucrose and starch content due to stress-induced biosynthesis enzyme deactivation, resulting in diminished seed size and weight (Zinta et al., 2018). Additionally, transcriptome analyses conducted on tobacco leaves under the combined stressors of heat and drought reveal the suppression of photosynthetic gene expression and the induction of genes associated with glycolysis and the pentose phosphate pathway (Rizhsky et al., 2002). Such molecular changes point towards reduced plant photosynthetic capacity and shortened seed filling duration, potentially influencing seed yield and quality. Seed quality is a critical component of marketability in soybean production, where the composition of seed protein, oil, and fatty acids is influenced by both genetic factors and the environment they grow. Although soybean-growing regions are prone to drought and heat stress, there is limited information on how soybean respond to these combined stresses (Jumrani and Bhatia, 2018; Ergo et al., 2018; Cohen et al., 2021b).

In addition, unique plant responses to combined stress have been identified to be governed by complex and distinct regulatory mechanisms (Zhang and Sonnewald, 2017; Cohen et al., 2021b; Sinha et al., 2023). A recent phenotypic-transcriptomic analysis of soybean plants subjected to individual and combined stressors revealed that different tissues displayed unique transcriptomic responses (Sinha et al., 2023). However, limited studies ascertained the impact of these synergistic stressors on physiology, spectral properties, yield, and quality attributes in soybean. On the other hand, previous studies used a few genotypes to understand how drought and heat stress impact plant performance. However, knowing how well high-yielding soybean cultivars handle these challenges is equally important, as they may exhibit more varied responses. Here, we report the impact of heat and drought stresses on ten soybeans’ genetic potential in terms of physiology, yield, and quality. This study also applies proximal sensing techniques to assess individuals and a combination of drought and heat.

## 2 MATERIALS AND METHODS

### 2.1 Plant materials

Ten different soybean cultivars belonging to maturity groups IV and V, which are commonly recommended for the Midsouth region, were used in this study. Among these cultivars, eight were commercially available for growers, and two (R15-2422 and R01-416F) were advanced breeding lines (Supplementary Table 1). R15-2422 was derived from crossing high-yielding conventional maturity group IV parents resistant to Cercospora leaf blight. R01-416F was the registered germplasm developed by crossing between Jackson and KS4896 to improve yield and nitrogen fixation under drought stress (Chen et al., 2007).

### 2.2 Crop husbandry

The experiment was conducted at Rodney Foil Plant Science Research Center of Mississippi State University, Mississippi, USA (33°28’ N, 88°47’ W) using a greenhouse facility. Four seeds per cultivar were sown in a 13.5L pot filled with farm soil. A 4 g of slow-release fertilizer Osmocote (N:P: K – 14:14:14, Hummert International) was added to the pot after sowing and top-dressed before flowering. A systemic insecticide Marathon 1% G (Imidacloprid, OHP, Mainland, PA) was applied to each pot (4 g) after seedling emergence to avoid infestation of sucking pests. After emergence, each pot was thinned down to a single plant. A total of 320 plants (ten cultivars × eight replicates × four treatments) were grown in a greenhouse under ideal conditions (32/24 °C day/night temperatures) for 50 days (until first flowering; R1stage, Fehr and Caviness, 1977). Plants were regularly monitored and watered through pre-programmed time-based drip irrigation to maintain moisture above 0.15 m^3^ m^-3^ volumetric water content (VWC).

### 2.3 Stress treatment conditions

At full bloom (R2 stage, Fehr & Caviness, 1977), the plants were divided into four treatments and moved to two greenhouses. One hundred and sixty pots were maintained in a greenhouse with 32 °C day temperature (current growing climate). Among them, 80 pots were provided with 100% irrigation (control, CNT), and others with 50% irrigation of the control (drought stress, DS). The remaining 160 pots were maintained in another greenhouse with 38 °C day temperature (warmer growing climate). Among them, 80 pots were provided with 100% irrigation (heat stress, HS), and the remaining with 50% irrigation (combined heat and drought, DS+HS). The nighttime temperatures were maintained at 24 °C in both the greenhouses. The thermostat, cooling pad, and ventail flaps were programmed to maintain the set temperature inside the greenhouse (Bheemanahalli et al., 2022b). HOBO data loggers (Onset Computer Corporation, Bourne, MA 02532, USA) were installed above the crop canopy for each treatment condition to monitor the microclimatic greenhouse conditions (temperature and relative humidity) throughout the experiment. Forty soil moisture probes (Model EM5b Soil Moisture, Decagon Devices, Inc., Pullman, WA, USA) periodically monitored soil moisture levels at 15 cm depth across all the treatments at 15-minute intervals. Stress was imposed for 30 days from the R2 (full bloom stage; Fehr and Caviness, 1977) to the R6 (full seed stage; Fehr and Caviness, 1977) stage. After 30 days of stress, all plants were grown under similar control and maintained until maturity.

### 2.4 Data collection

#### 2.4.1 Leaf pigments and physiological parameters

Leaf pigments such as the chlorophyll and anthocyanin indexes were recorded using a handheld Dualex® Scientific instrument (Force A DX16641, Paris, France). The physiological parameters (stomatal conductance and transpiration) were measured using a portable handheld LI600 porometer system integrated with a fluorometer (LI-COR Biosciences, Lincoln, USA) across the treatments. These parameters were recorded after every two-day interval throughout the stress period. At 14 days of stress, the photosynthesis was measured using portable LICOR 6800 (LI-COR Biosciences, Lincoln, USA) at 1500 μmol photons m^−2^ s^−1^, 420 μmol CO_2_ mol^−1^ air, and a constant flow rate of 600 μmol m^−2^ s^−1^, as recommended by the manufacturer. All the pigments and physiological parameters were measured from the third fully expanded trifoliate leaf from the apical end of the replicate plants. The canopy temperature was measured during solar noon using the handheld MI-2300 infrared radiometer (Apogee Instruments Inc., Logan, UT, USA) over the plant canopy.

#### 2.4.2 Leaf spectral signatures and vegetation indices

To evaluate the impact of treatments on leaf biophysical properties, leaf hyperspectral data (350-2500 nm) were collected using a PSR+3500 spectroradiometer (Spectral Evolution, Massachusetts, USA) on the third fully expanded trifoliate leaf between 10:00 h and 13:00 h during solar noon. The spectroradiometer was connected to a fiber optic cable and an internal light source. Four random replicates of each cultivar under each treatment were scanned thrice to reduce measurement noise. A white reference panelboard within the leaf clip calibrated the instrument every 30 minutes. Five sets of spectral bands were employed to calculate the vegetation indices (VIs) to match the proximal sensing to the commercially available MicaSense RedEdge multispectral sensor (Bheemanahalli et al., 2022b). These bands encompassed blue (centered at 475 nm/ bandwidth of 32), green (centered at 560 nm/ bandwidth of 28), red (centered at 668 nm/ bandwidth of 16), red-edge (centered at 717 nm/ bandwidth of 12), and near-infrared band (centered at 842 nm/ bandwidth of 85 nm), from which six VIs were derived: Chlorophyll index of green (CIgreen), chlorophyll index of red-edge (CIred-edge), chlorophyll vegetation index (CVI), normalized difference red-edge index (NDRE), Transformed Chlorophyll Absorption In Reflectance Index (TCARI), and Visible Atmospherically Resistant Index (VARI), using equations given in Supplementary Table 2.

#### 2.4.3 Yield and quality components

The replicated plants were manually harvested at physiological maturity (R8 stage; Fehr and Caviness, 1977) to obtain the yield and yield components. The shoot and pods were separated from each plant. The pods were counted and weighed before oven-drying them at 35 °C for 24 h and later threshed manually to obtain the seed weight. The number of seeds per plant was determined using a seed counter (NP5056-Model 850-2, LI-COR, Lincoln, NE, USA). The 100-seed weight was determined to estimate the impact of treatment on seed size. After collecting the yield, we assessed the soybean seed quality using a Perten DA7250 (Perten Instruments, Springfield, IL, USA). The quality of the seed sample was measured twice per replicate to ensure accurate results. The scanning was performed using the default setting and calibrations developed by the DA7250 manufacturer for soybean seed samples (Bheemanahalli et al., 2022a).

#### 2.4.4 RNA extraction and quantitative real-time polymerase chain reaction (qRT-PCR)

The leaves of all cultivars were sampled during the flowering and seed-setting stage, with four biological replicates for each treatment. Leaf samples weighing 200mg were ground with 1ml TRIzol using a pestle and mortar. Total RNA was isolated following the TRIzol manufacturer’s instructions (Invitrogen) and quantified using a BioPhotometer Plus (Eppendorf AG, Hamburg, Germany). To remove DNA contamination, 10ug of RNA was subjected to sequential DNase treatment using the TURBO DNA-free™ Kit (Ambion). Following the manufacturer’s instructions, reverse transcription was performed using 3ug RNA and the SuperScript IV reverse transcriptase first-strand synthesis kit (Invitrogen). PCR programs were run on an Applied Biosystems 96-Well Thermal Cycler (Eppendorf AG, Hamburg, Germany) for the DNase treatment and reverse transcription reaction. qRT-PCR was performed on an ABI 7500 Fast PCR System (ThermoFisher Scientific, Waltham, MA, USA), using the 2X PowerUp SYBR green master mix (Applied Biosystems, ThermoFisher Scientific) with the following settings: 50 °C for 2 min and 95 °C for 10 min followed by 40 cycles of 95 °C for 15 sec, 55 °C for 15 sec and 72 °C for 1 min. Gene expression analysis was carried out using three stress-responsive genes, namely GLYMA.10G23600, GLYMA.07G109100, and GLYMA.03G30040, based on homolog searches and literature (Wang et al., 2018; Xu et al., 2019). These genes correspond to drought, heat stress, and their combinations. Primer sequences can be found in Supplementary Table 3.

### 2.5 Statistical Analysis

The experimental design was a split-plot randomized complete block design, with treatment as the main plot factor and the cultivars as the subplot factors. The significance of treatment, cultivar, and their interaction for all the parameters was analyzed using the library “lmer”, “lsmeans” and “agricolae” (Bates et al., 2015; Lenth, 2016; Mendiburu and Yaseen, 2020). The post hoc Fisher’s Least Significant Difference (LSD) was used for mean separation. Differences were considered significant when *p*< 0.05. Data were analyzed using the statistical software R version 4.2.2 (https://www.R-project.org/, R Core). Correlation analyses were conducted to determine whether soybean cultivars exhibited unique responses to individual and combined stress treatments. The stress tolerance index (STI) was calculated for the ten soybean cultivars under drought, heat and combined treatments for physiology (chlorophyll content, anthocyanin, stomatal conductance, transpiration, canopy temperature, photosynthesis), leaf reflectance (CI green, CI red-edge, CVI, NDRE, TCARI, VARI), yield (pod number/weight, and seed number/ weight), and quality (protein, oil, linoleic acid, linolenic acid, oleic acid, sucrose) parameters using the formula defined by Fernandez, 1992.

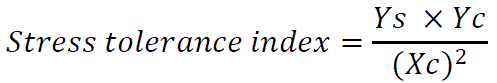

Ys is the phenotypic mean of a given cultivar under a given stress, and Yc is the phenotypic mean of a given cultivar under control. Xc is the mean yield of all cultivars under control. The cultivars were ranked based on the stress tolerance index value. A score from 1 (sensitive) to 10 (tolerant) was assigned to each cultivar based on the physiology, leaf reflectance, yield, and seed quality stress tolerance index. The library “ggpubr” was used for the bubble plot. All graphs were generated using the library “ggplot2” in R and Sigma Plot 14.5 (Systat Software, San Jose, CA, USA).

## 3 RESULTS

### 3.1 Temperature and soil moisture content

Based on the historical and projected monthly precipitation and maximum temperature during the reproductive period in the southern US, two soybean growing environments (current and warmer growing temperatures with optimum and low precipitation) were replicated during the flowering and seed-setting stages. Ten soybean cultivars were exposed to four treatments during flowering to the seed-setting stage (Figure 1). The volumetric water content was 0.15 m^3^ m^-3^ ± 0.03 under control, 0.06 m^3^ m^-3^ ± 0.01 under drought, 0.14 m^3^ m^-3^ ± 0.03 under heat, and 0.05 m^3^ m^-3^ ± 0.01 under combined stress for 30 days, during R1-R6 stages (Figure 1a). The average maximum daytime air temperature was maintained at 34.7 °C ± 1.8 under control and drought stress, while under heat and combined stress, the temperature was 8.4 °C higher than the control during the stress period (Figure 1b).

**Figure 1.**
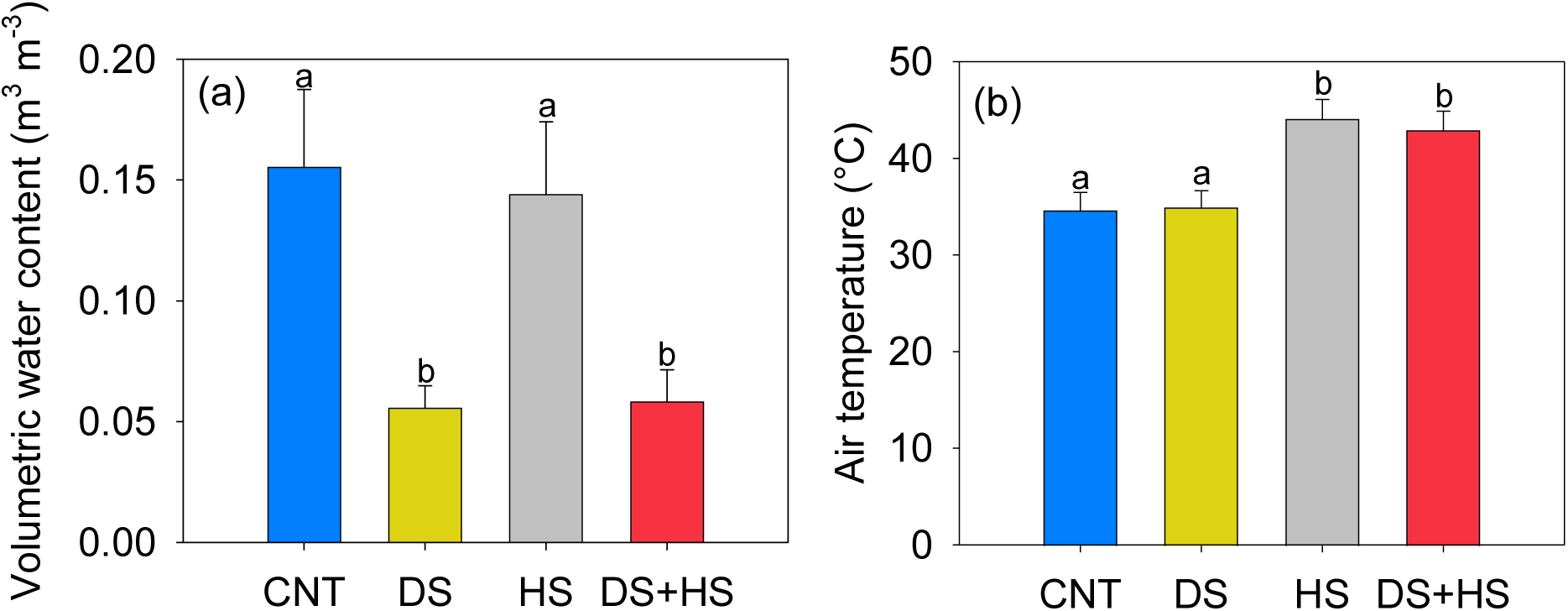
Volumetric water content. (A) and maximum air temperature (B) recorded during the experiment. The bar graphs represent the average over 30 days ± SD. CNT – control, DS – drought stress, HS – heat stress, and DS+HS – drought and heat stress. Means followed by the same letter are not significantly different at *p*< 0.05.

### 3.2 Pigments and physiological traits

Stress induced a significant (*p<* 0.001) effect on pigments and physiological parameters (Table 1). In contrast, the cultivar × treatment interaction effect was found to be significant (*p<* 0.05) for stomatal conductance (*gs*), transpiration rate, and anthocyanin content on 14 days of stress (R4 stage-full pod) (Table 1). Under drought stress, there was an initial increase in chlorophyll content (Supplementary Figure S1a), and it remained relatively stable under control and drought stress conditions. However, there was a 16% and 10% reduction under heat and combined stress, respectively, compared to the control (Table 1; Figure 2a). The cultivar DM45X61 (23% decrease) and LS5009XS (18% decrease) displayed a maximum reduction in chlorophyll content under heat and combined stress compared to the control (Figure 2a). R15-2422 performed consistently better under drought, heat, and combined stress, with the lowest reduction in chlorophyll content compared to the control. The interactive heat and drought stress led to a substantial increase (20%) in anthocyanin content (Figure 2b), with R15-2422 having a maximum increase (31%) among all the cultivars, compared to the corresponding control treatment (Figure 2b).

**Figure 2.**
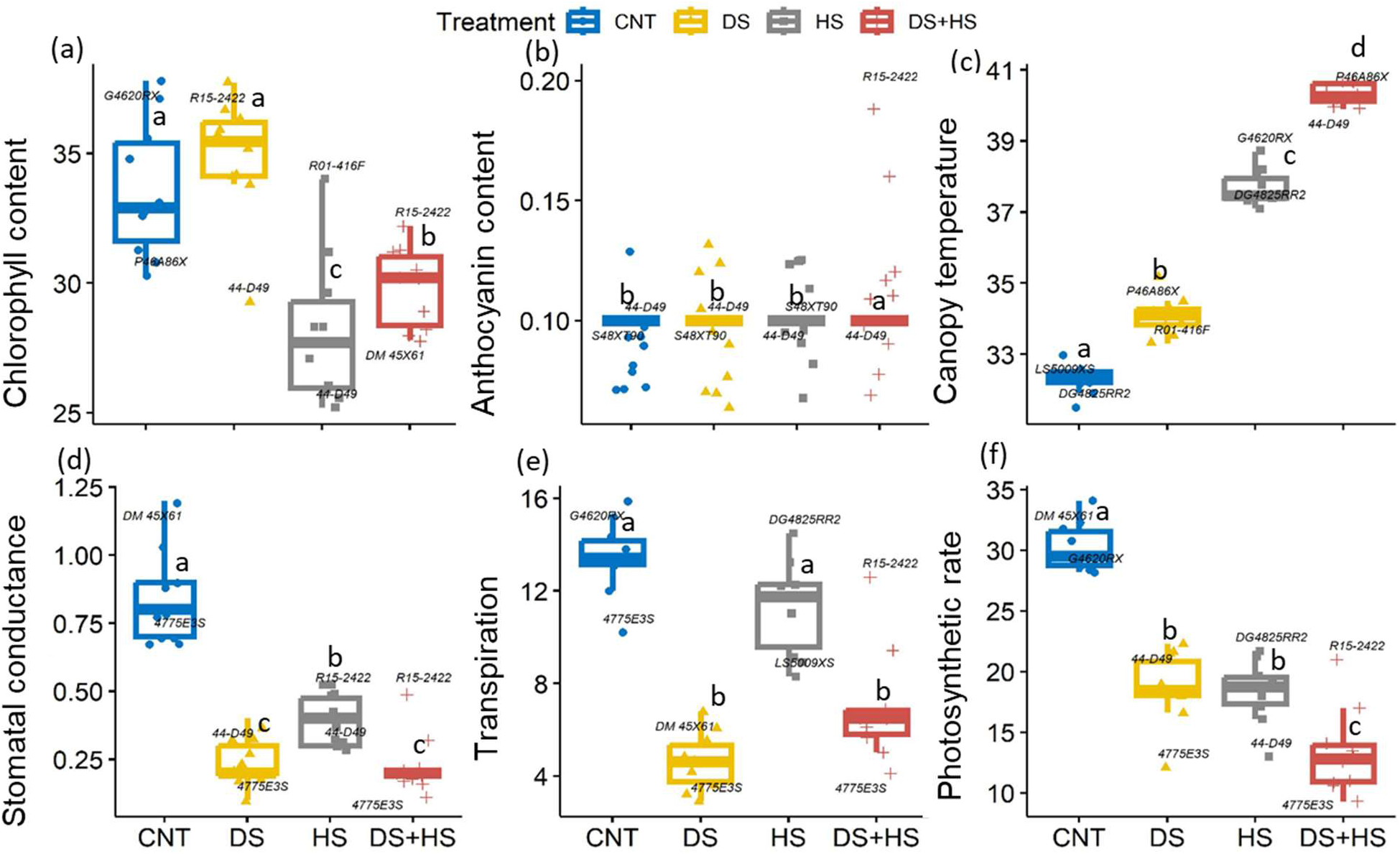
Effects of drought, heat, and their combination during the flowering seed setting stage on chlorophyll content. (µg cm^-2^, a), anthocyanin index (b), canopy temperature (°C, c), stomatal conductance (mol m^-2^ s^-1^, d), transpiration (mmol m^-2^ s^-1^, e), and photosynthetic rate (µmol m^-2^ s^-1^, f). CNT – control, DS – drought stress, HS – heat stress, and DS+HS – combined heat and drought stresses. Means followed by the same letter are not significantly different by the least significant difference (LSD) test at *p<* 0.05.

**Table 1.** Analysis of variance and mean values of the pigment, physiological, yield, and quality components of ten soybean cultivars (C) under control (CNT), drought (DS), heat (HS), and combined heat and drought (DS+HS) treatments (T).

| Trait | T | C | T x C | CNT | DS | HS | DS+HS |
| --- | --- | --- | --- | --- | --- | --- | --- |
| Chlorophyll content (Chl, $\mu\text{g cm}^{-2}$) | *** | *** | ns | 33.6<sup>a</sup> | 34.8<sup>a</sup> | 28.11<sup>b</sup> | 29.83<sup>b</sup> |
| Anthocyanin index (Anth) | ** | *** | * | 0.12<sup>b</sup> | 0.12<sup>b</sup> | 0.13<sup>b</sup> | 0.14<sup>a</sup> |
| Stomatal conductance (gs, $\text{mol m}^{-2} \text{s}^{-1}$) | *** | *** | *** | 1.02<sup>a</sup> | 0.38<sup>b</sup> | 0.92<sup>a</sup> | 0.06<sup>c</sup> |
| Transpiration rate (E, $\text{mmol m}^{-2} \text{s}^{-1}$) | *** | *** | *** | 8.29<sup>b</sup> | 5.51<sup>c</sup> | 11.70<sup>a</sup> | 2.67<sup>d</sup> |
| Canopy temperature (CT, $^{\circ}\text{C}$) | *** | * | ns | 32.0<sup>b</sup> | 34.0<sup>c</sup> | 37.6<sup>b</sup> | 40.3<sup>a</sup> |
| Photosynthetic rate (A, $\mu\text{mol m}^{-2} \text{s}^{-1}$) | *** | ns | ns | 29.6<sup>a</sup> | 19.0<sup>b</sup> | 18.3<sup>b</sup> | 13.1<sup>c</sup> |
| Pod number (PN, $\text{plant}^{-1}$) | *** | *** | * | 163<sup>a</sup> | 85.0<sup>c</sup> | 124.5<sup>b</sup> | 86.7<sup>c</sup> |
| Pod weight (PWt., $\text{g plant}^{-1}$) | *** | * | ns | 83.0<sup>a</sup> | 52.2<sup>c</sup> | 57.9<sup>b</sup> | 37.3<sup>d</sup> |
| Seed number (SN, $\text{plant}^{-1}$) | *** | *** | *** | 378.3<sup>a</sup> | 191.4<sup>c</sup> | 274.3<sup>b</sup> | 174.2<sup>d</sup> |
| Seed weight (SWt., $\text{g plant}^{-1}$) | *** | *** | *** | 55.9<sup>a</sup> | 36.0<sup>b</sup> | 36.9<sup>b</sup> | 23.8<sup>c</sup> |
| Hundred seed weight (HSWt., g) | *** | *** | ns | 14.8<sup>b</sup> | 18.9<sup>a</sup> | 13.7<sup>c</sup> | 13.9<sup>c</sup> |
| Protein (% dry basis) | *** | *** | ** | 40.3<sup>b</sup> | 43.3<sup>a</sup> | 40.6<sup>b</sup> | 40.7<sup>b</sup> |
| Oil (% dry basis) | *** | *** | *** | 23.2<sup>b</sup> | 20.5<sup>d</sup> | 23.8<sup>a</sup> | 22.8<sup>c</sup> |
| Linoleic acid (% dry basis) | *** | *** | * | 52.5<sup>a</sup> | 46.3<sup>b</sup> | 43.4<sup>c</sup> | 39.5<sup>d</sup> |
| Linolenic acid (% dry basis) | *** | *** | *** | 8.35<sup>c</sup> | 9.51<sup>a</sup> | 7.83<sup>d</sup> | 8.82<sup>b</sup> |
| Oleic acid (% dry basis) | *** | *** | ** | 22.7<sup>d</sup> | 28.2<sup>c</sup> | 33.5<sup>b</sup> | 36.4<sup>a</sup> |
| Sucrose (% dry basis) | *** | *** | ns | 4.56<sup>a</sup> | 4.49<sup>a</sup> | 4.14<sup>b</sup> | 4.52<sup>a</sup> |
| Chlorophyll index of green (CIgreen) | *** | *** | *** | 4.03<sup>a</sup> | 4.23<sup>a</sup> | 3.70<sup>b</sup> | 3.67<sup>b</sup> |
| Chlorophyll index of red-edge (CIred-edge) | *** | *** | *** | 0.93<sup>a</sup> | 0.95<sup>a</sup> | 0.80<sup>b</sup> | 0.77<sup>b</sup> |
| Chlorophyll vegetation index (CVI) | *** | *** | *** | 2.32<sup>b</sup> | 2.58<sup>a</sup> | 1.92<sup>c</sup> | 1.97<sup>c</sup> |
| Normalized Difference Rededge (NDRE) | *** | *** | *** | 0.31<sup>a</sup> | 0.32<sup>a</sup> | 0.28<sup>b</sup> | 0.28<sup>b</sup> |
| Transformed Chlorophyll Absorption in Reflectance Index (TCARI) | *** | *** | *** | 0.21<sup>a</sup> | 0.19<sup>b</sup> | 0.25<sup>a</sup> | 0.24<sup>a</sup> |
| Visible Atmospherically Resistant Index (VARI) | *** | *** | ** | 1.46<sup>b</sup> | 1.27<sup>c</sup> | 1.77<sup>a</sup> | 1.60<sup>a</sup> |

\*, \*\*, and \*\*\*, indicate significance levels at $p < 0.05$ , $p < 0.01$ , $p < 0.001$ , respectively. 'ns' indicates non-significance. Different letters in superscript indicate the significant treatment effect using the least significant difference (LSD) test at $p < 0.05$ .

There was a consistent declining trend in *gs* and transpiration rate parameters under drought and combined stress conditions (Figure 2). Heat and drought alone or combined increased canopy temperatures compared to the control. Cultivars exposed to combined stress had a higher canopy temperature (+8 °C ± 1.5) followed by heat (+5 °C ± 1.3) and drought stress (+2 °C ± 1.5) compared to the control (32 °C) (Figure 2c; Table 1). It was observed that an increase in canopy temperature had a negative association with the photosynthetic rate. Compared to control, the photosynthetic rate was reduced by 33% and 31% under drought and heat stress conditions, respectively (Figure 2f; Table 1). When both stresses were combined, the reduction in all physiological parameters resulted in a substantial drop (50%) in the photosynthetic rate (Figure 2). Different cultivars responded differently to individual and interactive stress. Specifically, the cultivar DM 45X61, which had the highest *gs* and photosynthetic rate under control, maintained a comparatively higher photosynthetic rate under drought. Under combined stress, R15-2422 recorded a maximum *gs*, transpiration, and photosynthetic rate, whereas 4775E3S showed the least performance. Under combined stress, R15-2422, with high chlorophyll content and stomatal conductance, maintained the highest photosynthetic rate compared to other cultivars (Figure 2). Despite significant reductions in stomatal conductance under drought and combined stress compared to heat, the rate of photosynthesis remained comparable under individual stresses and significantly reduced under combined stress. This result suggests that in addition to limited CO_2_ availability due to stomatal closure, the canopy temperature increase, and leaf pigment changes can further reduce photosynthesis in plants subjected to combined stresses.

### 3.3 Leaf spectral properties

To determine whether individual or combined stress induces different effects on plant health, leaf reflectance properties of all cultivars were measured 14 days after stress initiation. The VIs such as CIgreen, CIred-edge, CVI, and NDRE demonstrated significant variation between the cultivars (*p<* 0.001), treatments (*p<* 0.001), and cultivars × treatments (*p<* 0.01) interaction (Table 1). Under the heat and combined stress, CIgreen, CIred-edge, CVI, and NDRE significantly reduced by 12%, 15%, 15%, and 11% compared to the control, respectively (Table 2). Meanwhile, the VIs associated with the chlorophyll pigment did not vary significantly under drought compared to control. In contrast, TCARI and VARI decreased under drought stress (9% and 14%, respectively) and increased under individual heat (21%) and combined stress (18% and 10%, respectively) (Table 2). R01-416F and R15-2422 had the highest VI values under stress conditions, whereas DM45X61, DG4825RR2, and 44-D49 showed the least.

**Table 2.** Variation in leaf reflectance parameters (VIs) of soybean cultivars grown under control (CNT), drought (DS), heat (HS), and combined heat and drought (DS+HS) treatments.

| VIs | Cultivar | 44-D49 | 4775E3S | DG4825RR2 | DM 45X61 | G4620RX | LS5009XS | P46A86X | R01-416F | R15-2422 | S48XT90 |
| --- | --- | --- | --- | --- | --- | --- | --- | --- | --- | --- | --- |
| **CI green** | CNT | 3.67±0.17<sup>a</sup> | 3.77±0.17<sup>ab</sup> | 3.65±0.17<sup>b</sup> | 3.68±0.17<sup>a</sup> | 4.27±0.17<sup>a</sup> | 4.11±0.16<sup>a</sup> | 3.94±0.11<sup>a</sup> | 5.38±0.15<sup>a</sup> | 4.07±0.16<sup>a</sup> | 3.75±0.11<sup>b</sup> |
|  | DS | 3.65±0.1<sup>a</sup> | 3.98±0.1<sup>a</sup> | 4.13±0.1<sup>a</sup> | 4±0.1<sup>a</sup> | 4.48±0.1<sup>a</sup> | 4.35±0.15<sup>a</sup> | 4.05±0.12<sup>a</sup> | 4.84±0.16<sup>ab</sup> | 4.29±0.15<sup>a</sup> | 4.47±0.18<sup>a</sup> |
|  | HS | 3.23±0.14<sup>a</sup> | 3.36±0.14<sup>c</sup> | 2.94±0.14<sup>c</sup> | 2.85±0.14<sup>c</sup> | 3.52±0.1<sup>b</sup> | 3.98±0.32<sup>ab</sup> | 3.67±0.19<sup>ab</sup> | 4.45±0.25<sup>b</sup> | 3.85±0.32<sup>a</sup> | 3.55±0.12<sup>b</sup> |
|  | DS+HS | 3.47±0.21<sup>a</sup> | 3.46±0.21<sup>bc</sup> | 3.02±0.21<sup>c</sup> | 3.28±0.21<sup>bc</sup> | 3.58±0.21<sup>b</sup> | 3.49±0.18<sup>b</sup> | 3.58±0.11<sup>b</sup> | 3.61±0.2<sup>c</sup> | 4.05±0.18<sup>a</sup> | 3.6±0.14<sup>b</sup> |
| **CI red-edge** | CNT | 0.86±0.04<sup>a</sup> | 0.91±0.04<sup>a</sup> | 0.85±0.04<sup>a</sup> | 0.88±0.04<sup>a</sup> | 1.01±0.04<sup>a</sup> | 0.99±0.04<sup>a</sup> | 0.92±0.02<sup>a</sup> | 1.18±0.02<sup>a</sup> | 0.91±0.04<sup>a</sup> | 0.85±0.02<sup>b</sup> |
|  | DS | 0.81±0.01<sup>ab</sup> | 0.9±0.01<sup>a</sup> | 0.93±0.01<sup>a</sup> | 0.9±0.01<sup>a</sup> | 1.02±0.01<sup>a</sup> | 0.97±0.02<sup>ab</sup> | 0.95±0.03<sup>a</sup> | 1.07±0.04<sup>b</sup> | 0.98±0.02<sup>a</sup> | 1.02±0.04<sup>a</sup> |
|  | HS | 0.73±0.02<sup>c</sup> | 0.77±0.02<sup>b</sup> | 0.69±0.02<sup>b</sup> | 0.66±0.02<sup>b</sup> | 0.8±0.02<sup>b</sup> | 0.9±0.06<sup>b</sup> | 0.81±0.04<sup>b</sup> | 1±0.05<sup>b</sup> | 0.86±0.06<sup>a</sup> | 0.8±0.02<sup>b</sup> |
|  | DS+HS | 0.77±0.04<sup>bc</sup> | 0.77±0.04<sup>b</sup> | 0.69±0.04<sup>b</sup> | 0.73±0.04<sup>b</sup> | 0.8±0.04<sup>b</sup> | 0.76±0.03<sup>c</sup> | 0.8±0.02<sup>b</sup> | 0.77±0.03<sup>c</sup> | 0.88±0.03<sup>a</sup> | 0.79±0.02<sup>b</sup> |
| **CVI** | CNT | 2.03±0.14<sup>a</sup> | 2.25±0.08<sup>a</sup> | 1.93±0.22<sup>b</sup> | 2±0.11<sup>b</sup> | 2.7±0.11<sup>a</sup> | 2.43±0.09<sup>ab</sup> | 2.26±0.09<sup>ab</sup> | 3.33±0.19<sup>a</sup> | 2.28±0.19<sup>b</sup> | 1.98±0.08<sup>b</sup> |
|  | DS | 2.01±0.05<sup>a</sup> | 2.36±0.11<sup>a</sup> | 2.35±0.13<sup>a</sup> | 2.32±0.07<sup>a</sup> | 2.86±0.16<sup>a</sup> | 2.71±0.17<sup>a</sup> | 2.53±0.14<sup>a</sup> | 3.22±0.19<sup>a</sup> | 2.74±0.15<sup>a</sup> | 2.76±0.14<sup>a</sup> |
|  | HS | 1.62±0.07<sup>b</sup> | 1.78±0.12<sup>b</sup> | 1.37±0.11<sup>c</sup> | 1.36±0.11<sup>c</sup> | 1.95±0.19<sup>b</sup> | 2.3±0.16<sup>b</sup> | 2.06±0.16<sup>b</sup> | 2.54±0.15<sup>b</sup> | 2.25±0.25<sup>c</sup> | 2.03±0.1<sup>b</sup> |
|  | DS+HS | 1.82±0.14<sup>ab</sup> | 1.9±0.1<sup>b</sup> | 1.67±0.05<sup>bc</sup> | 1.75±0.14<sup>b</sup> | 2.08±0.16<sup>b</sup> | 2.1±0.13<sup>b</sup> | 2.08±0.09<sup>b</sup> | 2.09±0.15<sup>c</sup> | 2.26±0.12<sup>bc</sup> | 2.03±0.1<sup>b</sup> |
| **NDRE** | CNT | 0.3±0.01<sup>a</sup> | 0.31±0.01<sup>a</sup> | 0.3±0.01<sup>a</sup> | 0.3±0.01<sup>a</sup> | 0.33±0.01<sup>a</sup> | 0.33±0.01<sup>a</sup> | 0.31±0<sup>a</sup> | 0.37±0<sup>a</sup> | 0.31±0.01<sup>ab</sup> | 0.3±0.01<sup>b</sup> |
|  | DS | 0.29±0<sup>ab</sup> | 0.31±0<sup>a</sup> | 0.32±0<sup>a</sup> | 0.31±0<sup>a</sup> | 0.34±0<sup>a</sup> | 0.33±0.01<sup>ab</sup> | 0.32±0.01<sup>a</sup> | 0.35±0.01<sup>ab</sup> | 0.33±0.01<sup>a</sup> | 0.34±0.01<sup>a</sup> |
|  | HS | 0.27±0.01<sup>c</sup> | 0.28±0.01<sup>b</sup> | 0.25±0.01<sup>b</sup> | 0.25±0.01<sup>b</sup> | 0.28±0.01<sup>b</sup> | 0.31±0.02<sup>b</sup> | 0.29±0.01<sup>b</sup> | 0.33±0.01<sup>b</sup> | 0.3±0.02<sup>b</sup> | 0.28±0.01<sup>b</sup> |
|  | DS+HS | 0.28±0.01<sup>bc</sup> | 0.28±0.01<sup>b</sup> | 0.26±0.01<sup>b</sup> | 0.26±0.01<sup>b</sup> | 0.28±0.01<sup>b</sup> | 0.27±0.01<sup>c</sup> | 0.28±0<sup>b</sup> | 0.28±0.01<sup>c</sup> | 0.31±0.01<sup>ab</sup> | 0.28±0.01<sup>b</sup> |
| **TCARI** | CNT | 0.22±0.02<sup>b</sup> | 0.2±0.02<sup>b</sup> | 0.25±0.02<sup>b</sup> | 0.21±0.02<sup>bc</sup> | 0.17±0.02<sup>b</sup> | 0.18±0.01<sup>b</sup> | 0.2±0.01<sup>ab</sup> | 0.13±0.01<sup>c</sup> | 0.2±0.01<sup>a</sup> | 0.22±0.01<sup>a</sup> |
|  | DS | 0.22±0.01<sup>b</sup> | 0.2±0.01<sup>b</sup> | 0.19±0.01<sup>c</sup> | 0.19±0.01<sup>c</sup> | 0.17±0.01<sup>b</sup> | 0.17±0.01<sup>b</sup> | 0.18±0.01<sup>b</sup> | 0.15±0.01<sup>c</sup> | 0.17±0.01<sup>a</sup> | 0.16±0.01<sup>b</sup> |
|  | HS | 0.26±0.01<sup>a</sup> | 0.25±0.01<sup>a</sup> | 0.31±0.01<sup>a</sup> | 0.31±0.01<sup>a</sup> | 0.24±0.01<sup>a</sup> | 0.19±0.03<sup>b</sup> | 0.22±0.02<sup>a</sup> | 0.18±0.01<sup>b</sup> | 0.22±0.03<sup>a</sup> | 0.21±0.01<sup>a</sup> |
|  | DS+HS | 0.24±0.02<sup>ab</sup> | 0.23±0.02<sup>a</sup> | 0.28±0.02<sup>ab</sup> | 0.26±0.02<sup>ab</sup> | 0.22±0.02<sup>a</sup> | 0.23±0.01<sup>b</sup> | 0.22±0.01<sup>a</sup> | 0.22±0.01<sup>a</sup> | 0.21±0.01<sup>a</sup> | 0.22±0.01<sup>a</sup> |
| **VARI** | CNT | 1.42±0.08<sup>b</sup> | 1.24±0.08<sup>b</sup> | 1.72±0.08<sup>b</sup> | 1.47±0.08<sup>b</sup> | 1.11±0.08<sup>b</sup> | 1.19±0.15<sup>ab</sup> | 1.38±0.04<sup>a</sup> | 1.09±0.11<sup>bc</sup> | 1.49±0.15<sup>a</sup> | 1.57±0.09<sup>a</sup> |
|  | DS | 1.42±0.05<sup>b</sup> | 1.26±0.05<sup>b</sup> | 1.35±0.05<sup>b</sup> | 1.29±0.05<sup>b</sup> | 1±0.05<sup>b</sup> | 1.14±0.05<sup>b</sup> | 1.12±0.06<sup>b</sup> | 0.95±0.07<sup>c</sup> | 1.1±0.05<sup>b</sup> | 1.14±0.08<sup>b</sup> |
|  | HS | 1.75±0.09<sup>a</sup> | 1.72±0.09<sup>a</sup> | 2.25±0.09<sup>a</sup> | 2.14±0.09<sup>a</sup> | 1.62±0.09<sup>a</sup> | 1.41±0.15<sup>a</sup> | 1.57±0.11<sup>a</sup> | 1.32±0.1<sup>ab</sup> | 1.49±0.15<sup>a</sup> | 1.48±0.08<sup>a</sup> |
|  | DS+HS | 1.68±0.13<sup>ab</sup> | 1.57±0.13<sup>a</sup> | 1.61±0.13<sup>b</sup> | 1.69±0.13<sup>b</sup> | 1.42±0.13<sup>a</sup> | 1.33±0.07<sup>ab</sup> | 1.43±0.09<sup>a</sup> | 1.38±0.08<sup>a</sup> | 1.43±0.07<sup>a</sup> | 1.48±0.09<sup>a</sup> |

Values represent mean (n = 10) ± SE for the leaf reflectance parameters (VIs). Different letters in superscript indicate the significant treatment effect for a given parameter between treatments. The mean values were separated using the least significant difference (LSD) test at $p < 0.05$ .

### 3.4 Yield parameters

The yield parameters showed significant variation between treatments (*p<* 0.001, Table 1). Not surprisingly, more pods were observed under control (163 plant^-1^), followed by heat stress (125 plant^-1^) compared to combined stress. The drought and combined stress impact on pod number were significantly *at par* with a reduction of ∼47% compared to control (Table 1). However, pod weight was reduced by 37% under drought, with the combined stress being the most severe (55% decrease) compared to control. Among the cultivars, the variability in percent reduction in seed yield under individual drought and heat stress ranged from 25 to 44% and 28 to 42%, whereas under interactive stress treatment, recorded 53 to 63% with maximum reduction displayed in 44-D49 and G4620RX compared to control (Figure 3). Significant seed number and weight reductions were observed under heat (27% and 34%) and drought (49% and 36%). The hundred seed weight was increased significantly by 28% under drought stress compared to control, whereas it was decreased by 8% under heat and 6% under the combined stress (Table 1, Figure 3). Even though the hundred seed weight under the combined stress was comparable (∼36 g) with heat stress, a significant reduction in seed number under the combination stress resulted in a pronounced decrease in seed yield (more than 50%) compared to control (Table 1, Figure 3). We observed more aborted and empty pods with small, wrinkled seeds under the heat and combined stress, whereas few but bigger seeds were found in the drought-stressed plants (Figure 3d).

**Figure 3.**
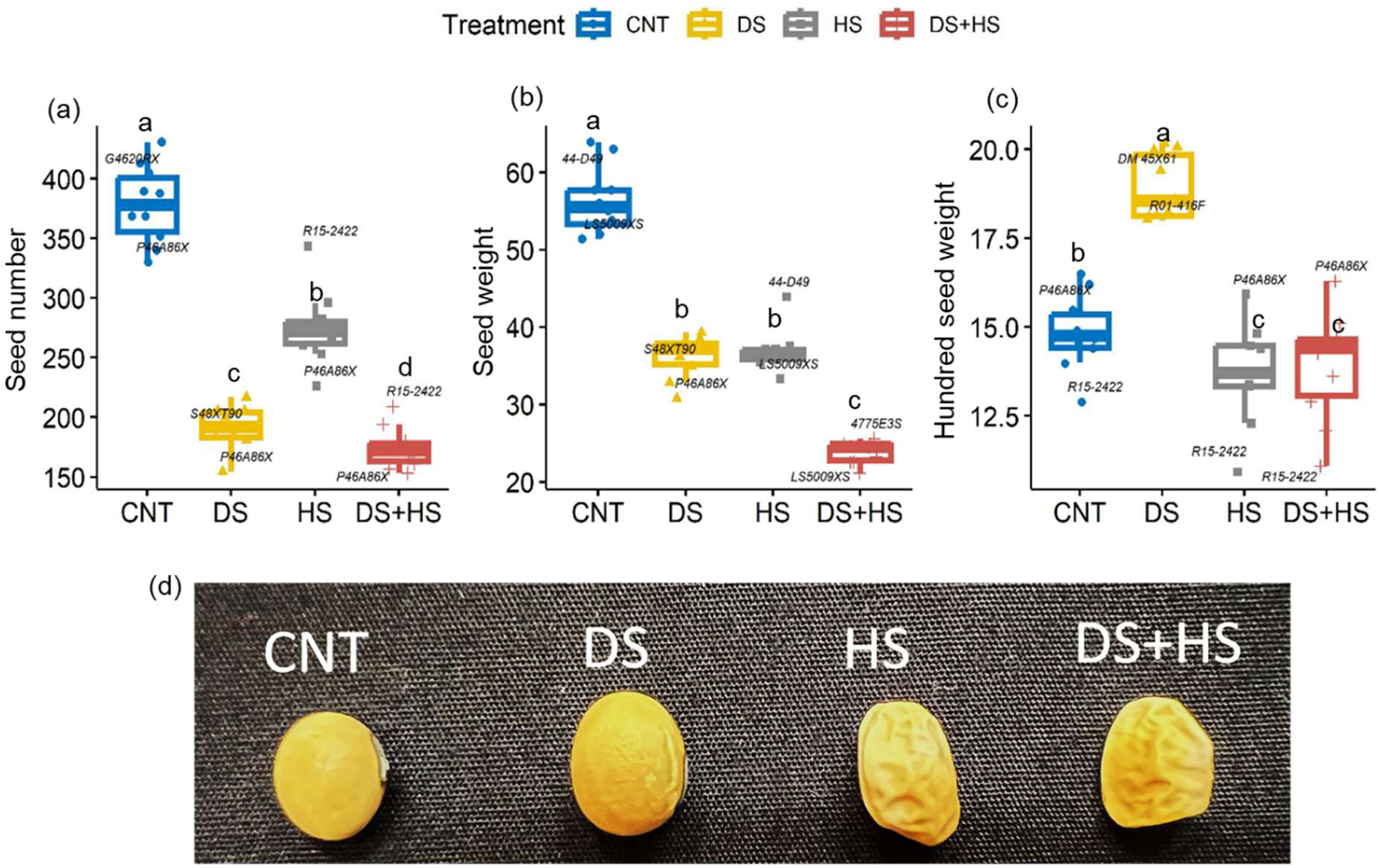
Effects of drought, heat, and combined heat and drought stresses on seed weight. (g plant^-1^, a), seed number (plant^-1^, b), and hundred seed weight (g, c). Pictorial representation of individual seed size under four treatment conditions (d). CNT – control, DS – drought stress, HS – heat stress, and DS+HS – combined heat and drought stresses. Means followed by the same letter are not significantly different by the least significant difference (LSD) test at *p<* 0.05.

### 3.5 Seed quality

Significant treatment (*p<* 0.001), cultivar (*p<* 0.001), and treatment × cultivar interaction (*p<* 0.01) were observed for protein and oil content (Table 1). All the cultivars responded differentially across the treatments for protein and oil content. Compared to the control, protein content increased by 8% under drought, while oil content decreased by 11%. The protein content was similar to control under individual heat and combined stress. Cultivar R15-2422 had a maximum protein content under drought, heat, and control conditions, and the minimum was observed in DG4825RR2 under individual stress conditions (Figure 4a). In contrast, the average oil content across the cultivars increased under heat, whereas under combined heat and drought stress, a reduction was observed, with an exception noticed in the cultivars; R15-2422, R01-416F, and DG4825RR2, and the least was in 44-D49 (Figure 4b).

**Figure 4.**
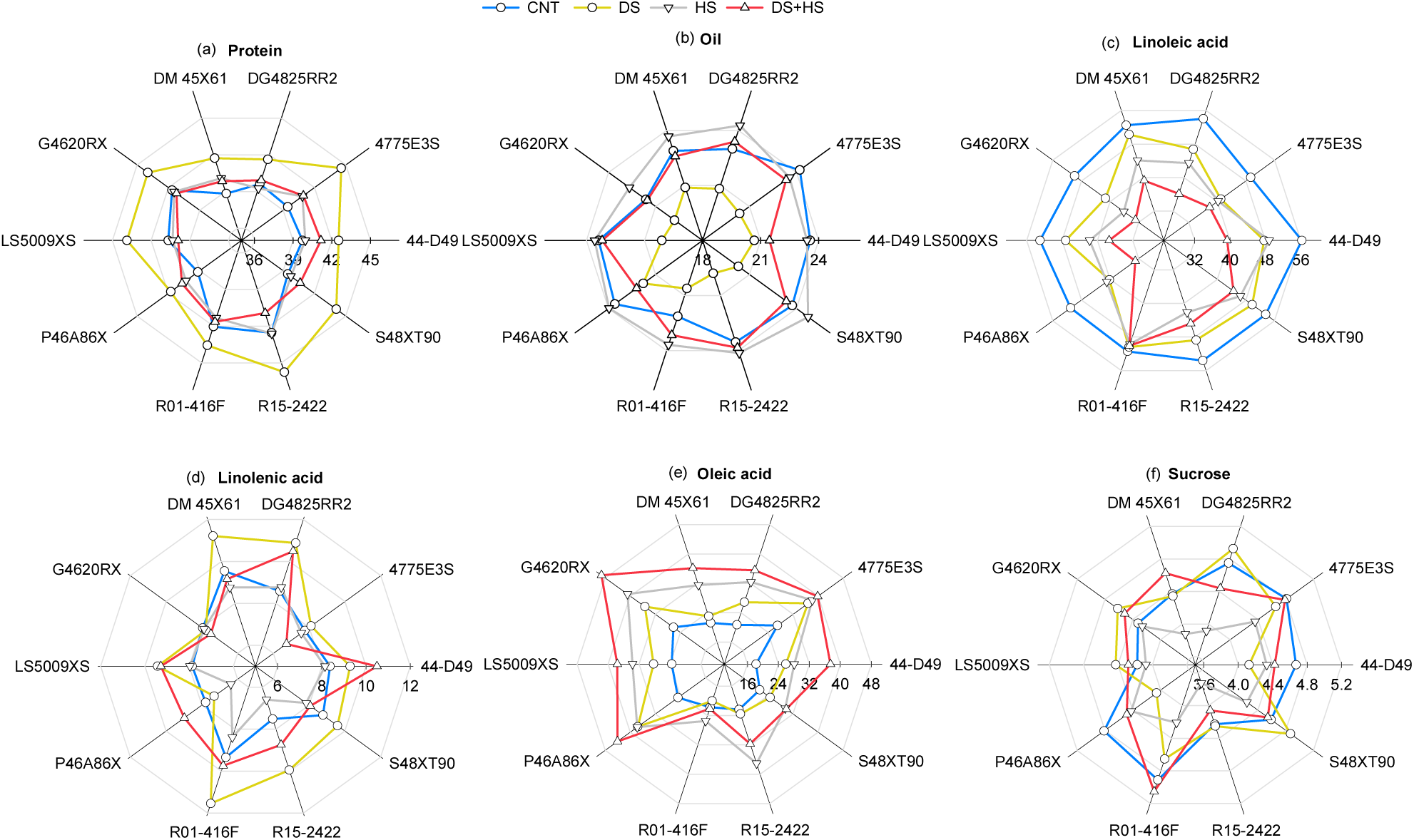
Effects of drought, heat, and combined stresses on protein. (% dry basis, a), oil (% dry basis, b), linoleic acid (% dry basis, c), linolenic acid (% dry basis, d), oleic acid (% dry basis, e), and sucrose (% dry basis, f). CNT – control, DS – drought stress, HS – heat stress, and DS+HS – combined heat and drought stresses.

**Figure 5.**
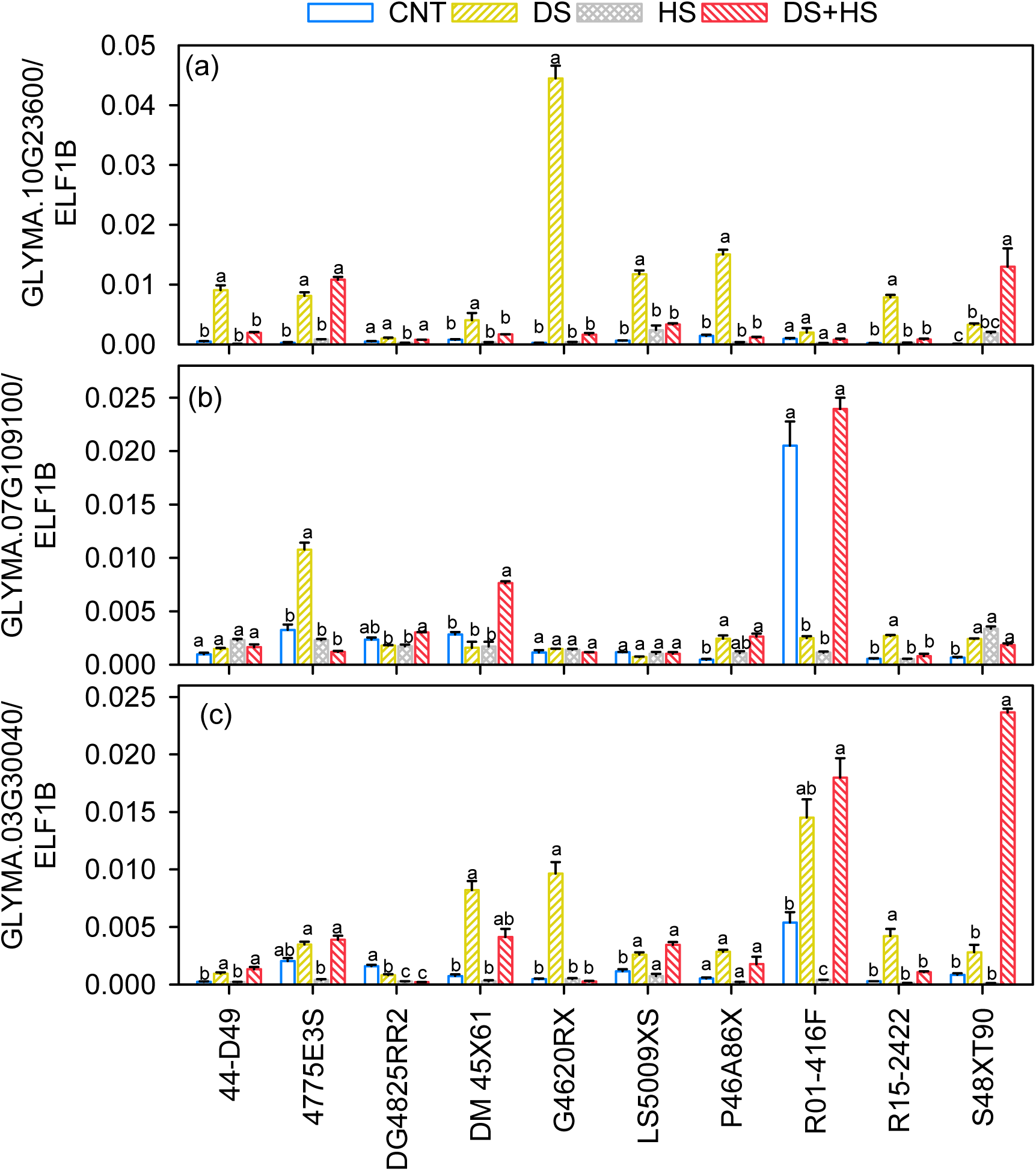
Quantitative Reverse Transcription Polymerase Chain Reaction. (qRT-PCR) of stress marker genes during soybean seed setting under various conditions. The test was conducted using stress-responsive genes, which include (a) the drought-responsive gene GLYMA.10G23600, (b) the heat-responsive gene GLYMA.07G109100, and (c) the combined stress-responsive gene GLYMA.03G30040. The data represent mean values and standard errors derived from four biological replicates. A cultivar with different letters indicates significant differences based on the Tukey HSD test at *p<* 0.05.

Soybeans are a rich source of linoleic and linolenic acids. However, when exposed to stressors, the composition of these essential fatty acids changed significantly. For example, linoleic acid, the most abundant unsaturated fatty acid in soybean, decreased under all stressors. The maximum reduction was recorded under combined stress (25%), while the minimum reduction occurred under drought (12%) (Figure 4c). Although soybeans contain lower amounts of linolenic acid than linoleic acid, there was a significant increase in linolenic acid under drought and combined stress compared to the control. On the other hand, the monounsaturated fatty acid, oleic acid, showed a higher accumulation under combined stress conditions followed by individual heat and drought stress (Table 1; Figure 4). Under individual and combined stress, the cultivars with the highest linoleic content, R01-416F, and R15-2422, exhibited a reduced level of oleic acid (Figure 4e). In contrast, the opposite is true, indicating a discernible trade-off between these fatty acids. The stress tolerance and yield of soybeans depend on this tradeoff and the interaction between protein and oil content.

### 3.6 Gene expression response of stress-responsive genes in soybean to stressors

GLYMA.07G109100 encodes a pentatricopeptide repeat-containing protein (PRR) that has a protective role in scavenging reactive oxygen species (ROS) specific to heat stress (Xu et al., 2019). However, less variation among the heat-stressed group was observed among ten cultivars, whereas drought-stressed 4775E3S, P46A86X, R15-2422, and combinational stress-treated DM45X61 displayed a significantly higher transcript level compared to control. The lack of significant changes across treatments may suggest the malfunction of oxidoreductase following long-term stress, resulting in a reduced ability to remove ROS. Additionally, GLYMA.03G30040, a homolog of AT5G06760, which encodes late embryogenesis abundant proteins, exhibited significant induction under individual drought and combinational stress conditions. Among the cultivars, gene expression was significantly induced under drought conditions compared to control groups for 44-D49, DM45X61, G4620RX, LS5009XS, and R15-2422. Additionally, S48XT90 exhibited a substantial induction of gene expression under combined stress conditions.

### 3.7 Stress tolerance index

The stress tolerance index (STI) was used to determine soybean cultivars’ tolerance to individual and combined stresses. The cultivar DM45X61 displayed higher tolerance, followed by 44-D49 and R15-2422 under heat and drought stress for the physiological parameters (Figure 6a). Under combined stress, the cultivar 44-D49 had a higher tolerance rank for the physiological parameters, followed by G4620RX and R15-2422 (Figure 6a). The cultivar R01-416F had higher tolerance in terms of leaf reflectance properties, consistently demonstrating higher quality and gene expression rankings across all three treatments, except for yield traits. For the yield parameters, cultivars 44-D49 were tolerant under drought and heat stress (Figure 6c). However, the cultivar DM45X61, which performed better in terms of physiology under individual heat and drought, did not show the highest rank for yield, possibly due to the negative impacts of the stress on the pollen viability and reproductive failure (Bheemanahalli et al., 2019; Poudel et al., 2023b). Meanwhile, under combined stress, the cultivar G4620RX, with the highest tolerance for physiological performance, showed better ranking for yield parameters (Figure 6a). Similar to R01-416F, 4775E3S also displayed a higher tolerance rank for the gene expression across the stress treatments (Figure 6e). Based on the average tolerance rank across the parameters, the cultivars R15-2422, G4620RX, and 44-D49 were the tolerant cultivars across the treatments. Similarly, the cultivar R01-416F, followed by 44-D49, had the highest stress tolerance rank for quality traits under the individual and combined heat and drought stress (Figure 6d).

**Figure 6.**
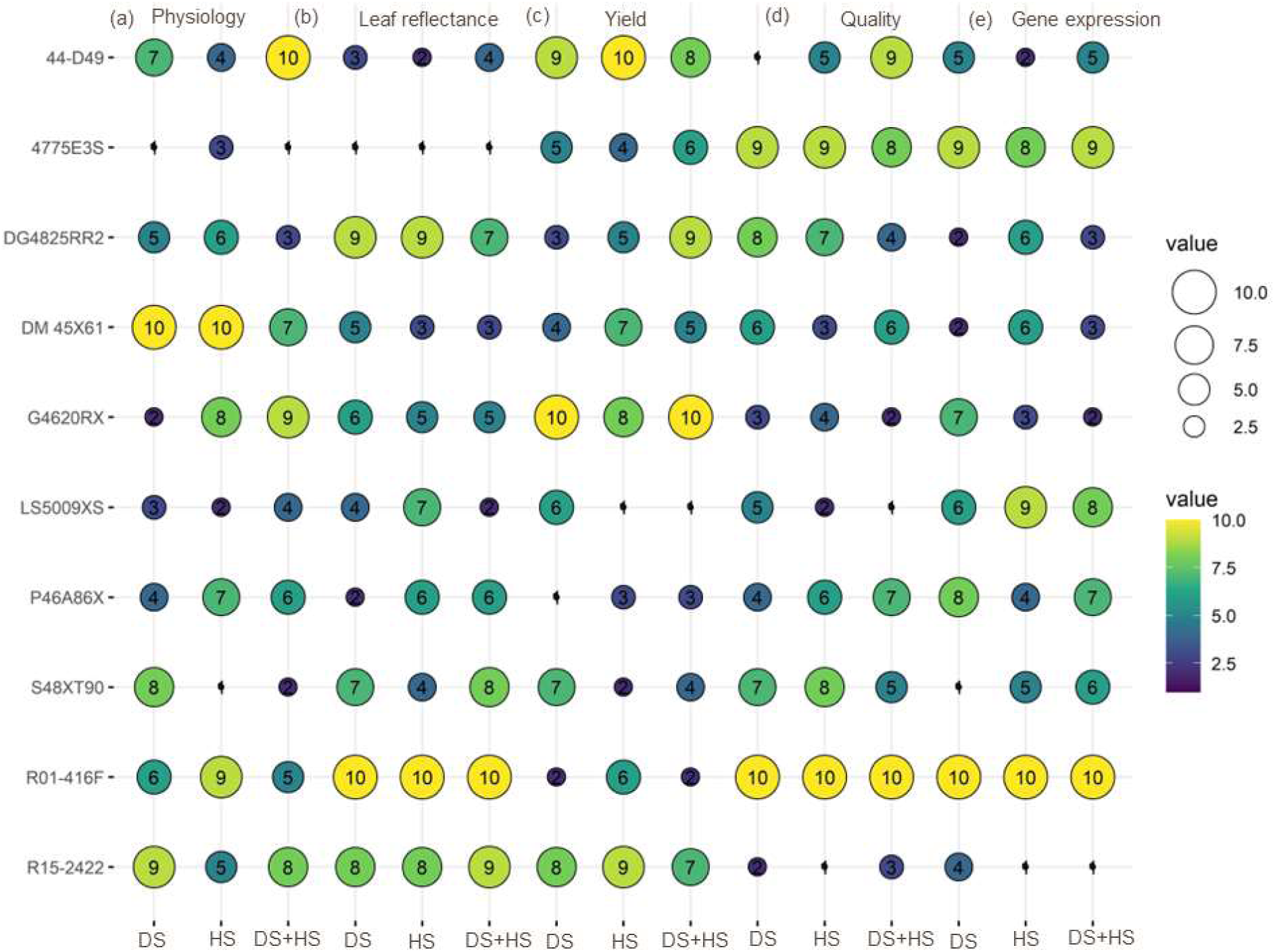
Bubble plot showing the average stress tolerance index values for physiology. (chl, anth, gs, E, CT, A; a), leaf reflectance (CI green, CI red-edge, CVI, NDRE, TCARI, VARI; b), yield (PN, PWt., SN, SWt., HSWt.; c), seed quality (protein, oil, linoleic acid, linolenic acid, oleic acid, sucrose; d), and gene expression (drought-responsive gene GLYMA.10G23600, heat-responsive gene GLYMA.07G109100, combined stress-responsive gene GLYMA.03G30040; e). A cultivar with a larger bubble size (yellow color) indicates higher stress tolerance and vice versa. DS – drought stress, HS – heat stress, and DS+HS – combined heat and drought stresses.

## 4 DISCUSSION

Paradigm shift occurring under combined drought and heat stress compared to the individual stressors is gaining prominence in soybean and other crops (Zandalinas et al., 2018; Ergo et al., 2018; Lawas et al., 2018; Zandalinas et al., 2020; Cohen et al., 2021a; b; Bheemanahalli et al., 2022b). Recent research shows that soybeans had a synergistic effect of combined heat and drought stresses, particularly affecting source-sink balance and yield (Du et al., 2023). The present results show that the interactive heat and drought in soybeans had a higher impact on physiology, yield, and seed composition compared to single stress at the cultivar level.

### 4.1 Interactive stress-induced changes in physiology and leaf reflectance properties in soybean

Drought stress significantly lowers stomatal conductance, reducing transpiration rate (Figure 2, 8). Notably, the drought-stressed plants-maintained greenness similar to the control (Figure 2a; Supplementary Figure S1a), by sustaining chlorophyll pigment content (Jurik, 1986). Likewise, heat-stressed plants increased their transpiration rate (Figure 2e) to maintain a cooler canopy (Sinha et al., 2022; Poudel et al., 2023a). As an adaptive strategy (differential transpiration) under interactive heat and drought, plants close leaf stomata while flower and pod stomata remain open (Cohen et al., 2021b; Sinha et al., 2022). Whereas comparatively higher transpiration was observed in the reproductive tissue than the vegetative as a protective mechanism to avoid overheating (Sinha et al., 2022; Vennam et al., 2023a). The canopy temperature, an integrative trait that reflects the plant water status, increased by 8 °C under combined stress had a pronounced disruption of photosynthesis and stomatal conductance, consistent with previous study (Cohen et al., 2021b). Combined stress resulted in a synergistic adverse effect on plant traits with a greater negative impact on the physiological traits than individual drought stress (Figure 2). The significant impact of treatment leaf reflectance properties indicates the changes in pigment accumulation, which can be used to determine plant health. The VIs associated with greenness, such as CI green, CI red edge, CVI, and NDRE, were comparable with manual measurements under stress (Lima et al., 2020; Aldubai et al., 2022). This also supports the hypothesis that stress-induced changes in the absorption, reflectance, and transmittance of radiation from the leaf vary with genetics (Walter-Shea et al., 1991; Kataria et al., 2014). As reported in other crops (Bheemanahalli et al., 2022b; Brewer et al., 2022), changes observed with the VIs unravel the application of proximal sensing to quantify the impact of stressors in soybeans during flowering. Differential response of VIs between control and treatments among cultivars indicates greater variability in stress tolerance (Figure 7).

**Figure 7.**
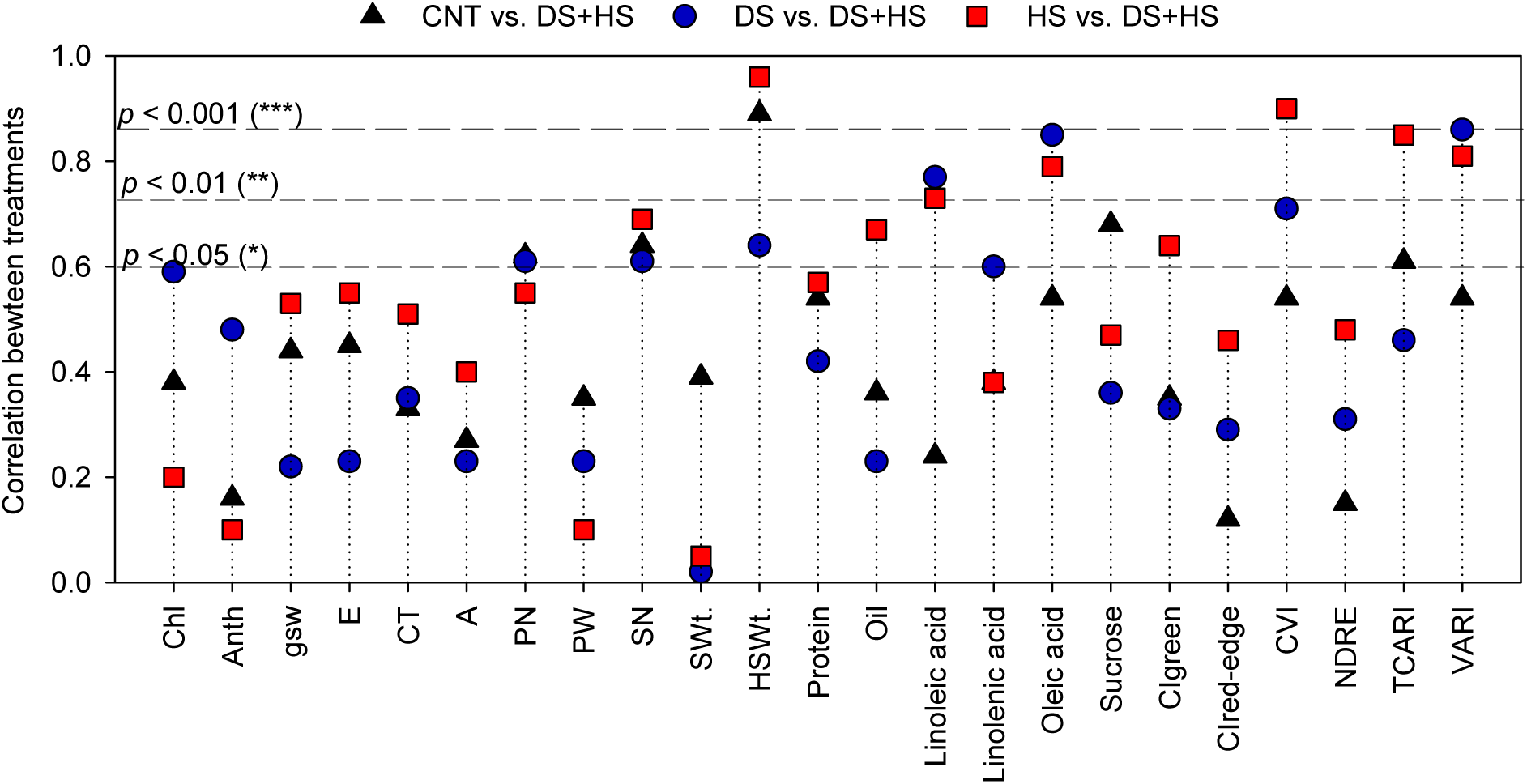
Correlation of traits between single and combined stress treatments. *, **, and ***, indicate significance levels at *p<* 0.05, *p<* 0.01, *p<* 0.001, respectively. CNT – control, DS – drought stress, HS – heat stress, and DS+HS – combined heat and drought stresses. \**p* < 0.05, \*\**p* < 0.01, \*\*\**p* < 0.001 indicate a significant correlation between treatment for a given trait. Traits acronyms are given in Table 1.

### 4.2 The implication of stress-responsive genes into phenotypic performance

The physiological and morphological changes due to various abiotic stresses are determined by genes that are governing in different hormonal signaling and regulatory pathways (Zhang et al., 2022; Kumar et al., 2023). Utilizing RT-qPCR, the expression patterns of well-studied classic stress-responsive genes were analyzed in response to drought, heat, and combinatorial treatments. Specifically, the drought-responsive marker GLYMA.10G23600 is a homolog of Arabidopsis RD29A and RD29B, two drought-inducible genes involved in the abscisic acid (ABA) signaling pathway, which plays a critical role in orchestrating plant responses to drought stress (Msanne et al., 2011). In this study, GLYMA.10G23600 showed significant induction in genotype G4620RX under drought conditions, correlating with its highest changes in stomatal conductance, transpiration rate, and chlorophyll content. The cultivar S48XAT90 exhibited significant gene induction under combined stress, which corresponds to noticeable physiological and morphological alterations. It has been implicated that ABA plays center stage in regulating stomatal movement, transpiration, and chlorophyll degradation (Hsu et al., 2021; Bharath et al., 2021). This correlation between GLYMA.10G23600 expression and phenotypes presented by G4620RX and S48XAT90 suggests a potential for enhanced stress response through modulated ABA signaling. Future research should focus on clarifying the key regulatory factors involved in ABA signaling in these two varieties.

Furthermore, the heat stress marker GLYMA.07G109100 encodes a pentatricopeptide repeat-containing protein (PRR), which is critical for ROS scavenging (Yu et al., 2021a). It displayed less variation in expression among heat-stressed groups. This observation raises the possibility that prolonged heat stress could produce excess ROS that exceeds cultivar’s capacities to maintain redox signaling (Fortunato et al., 2023). Nevertheless, cultivars 4775E3S, P46A86X, and R15-2422 demonstrated significant gene expression induction under drought, while DM45X61 only exhibited increased mRNA induction in response to combined heat and drought stresses. These patterns possibly indicate a well-maintained ROS scavenging among the above-mentioned varieties during long-run drought or combined stress.

In addition, the combined heat and drought stress marker GLYMA.03G30040 is homologous to Arabidopsis late embryogenesis abundant (LEA) 4-5. Most LEA proteins are considered a subset of hydrophilins with a specialized function in retaining water molecules (Battaglia et al., 2008). Previous RNA studies revealed significant induction of this particular stress marker during short-term drought, salt or combined drought and heat stress in soybeans (Wang et al., 2018; Guo et al., 2023). However, very few studies have characterized its expression patterns in soybeans exposed to long-term stressors. Interestingly, in the current study, the gene expression of GLYMA.03G30040 is significantly elevated in drought-stressed cultivars 44-D49, DM45X61, G4620RX, LS5009XS, and R15-2422 and combined stressed S48XT90. This line of evidence of GLYMA.03G30040 induction implies LEA could also accumulate during prolonged drought or combined stressors. Overall, the stress-responsive genes exhibit varying degrees of transcript alternations across varieties. Functional proteins or hormone pathways are implied to be involved in these changes, providing valuable insights into the molecular mechanisms underlying the distinct phenotypic performance of stressors.

### 4.3 Interactive stress-induced alterations in yield components

Plant yield is the complex integration of principal events, including physiological and biochemical processes, flowering phenology, and seed-filling rate (Figure 8). The negative impact of combined stress on yield was 2-fold higher compared to drought or heat alone (Figure 3), with a similar response recorded in other crops (Bheemanahalli et al., 2022b). On the other hand, despite significant difference in seed number between drought and heat, seed weight remains similar, indicating that water limitation has more damaging impacts on pod number than heat stress (Poudel et al., 2023a). With the increase in canopy temperature by 8 °C under the combined stress, there was more than a 50% reduction in seed weight and number. A significantly fewer number of pods (37%) and seeds (49%) were observed under drought stress (Figure 3c-d). However, upon rehydration, the remaining seeds were larger, which might be due to the translocation of photosynthates produced after rewatering to a limited number of seeds (source) (Ney et al., 1994; Poudel et al., 2023b). The tradeoff between the seed number and seed weight has been a well-documented adaptive strategy in crops (Griffiths et al., 2015; Cohen et al., 2021b). Heat stress alone or in combination with drought-induced early senescence and shortened seed-filling duration resulted in fewer pods with small, wrinkled seeds (Figure 3d; Jumrani and Bhatia, 2018; Pradhan et al., 2012). R15-2422 had the least reduction in seed number under heat and combined stress conditions compared to control due to better physiological traits (Figure 7). However, a significant decrease in hundred-seed weight made this cultivar highly susceptible to combined heat and drought stress (Figure 3c). This finding indicates poor assimilate translocation between source and sink. In addition, the higher number of empty pods and aborted seeds observed under heat and combined stress suggests the sensitivity of reproductive failure, as seen in other studies (Bheemanahalli et al., 2022b; Koti et al., 2005b; Sinha et al., 2022). Currently, growing soybean cultivars are bred for a higher yield. However, none of these cultivars can tolerate combined stress during reproductive and seed filling. Therefore, a particular focus should be given to developing cultivars with superior physiological traits that minimize reproductive failure to achieve higher yields under combined stress.

**Figure 8.**
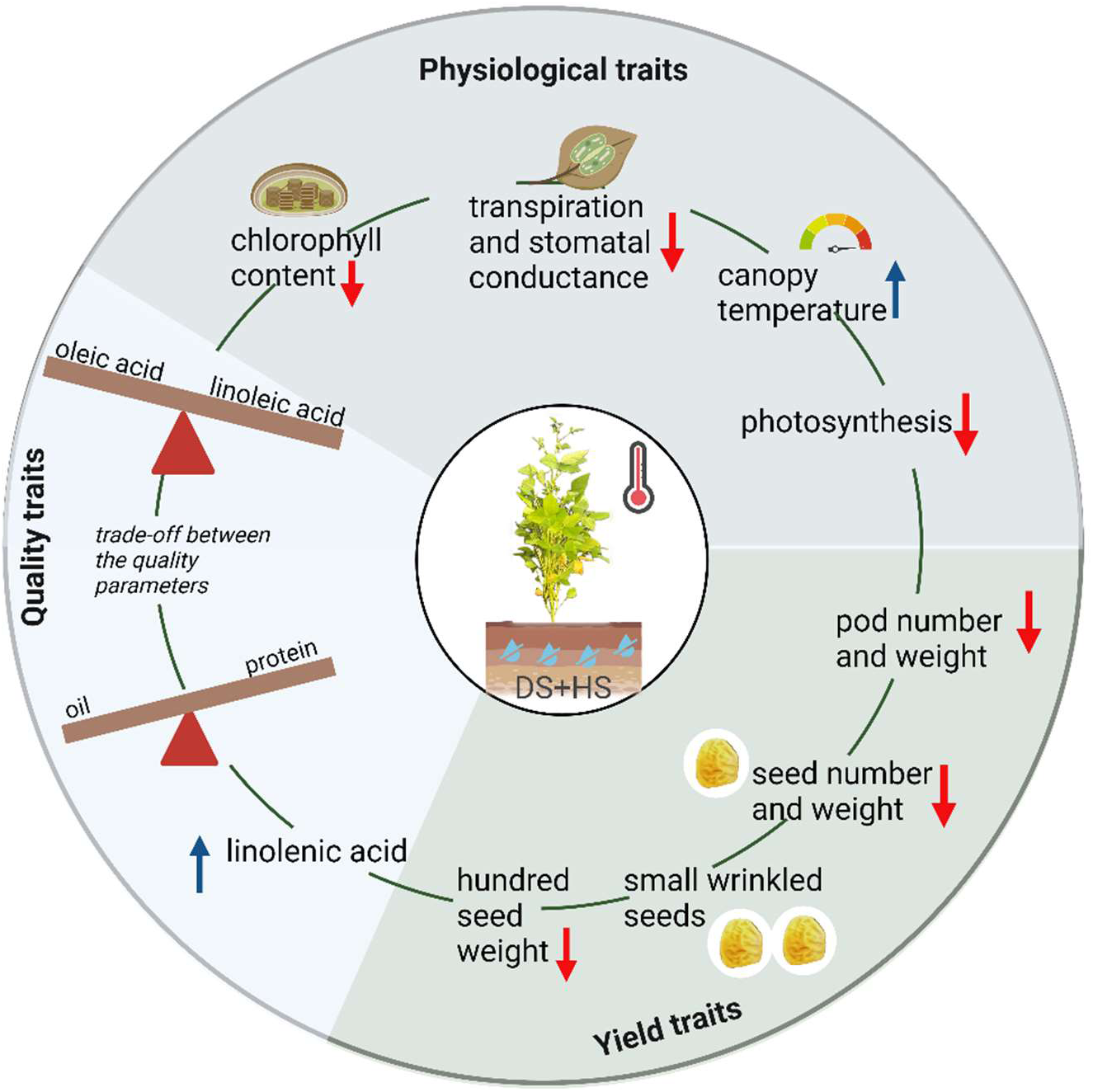
Summary of combined heat and drought stress. (DS+HS) impact on soybean morpho-physiology, yield, and quality traits in soybean. Illustration created using the Biorender.

### 4.4 Trade-off between yield and seed quality components

Protein and oil content of soybean showed an inverse correlation across treatments, with a stronger negative correlation under drought (r = –0.81, *p*< 0.01), consistent with other studies (Bellaloui et al., 2015; Mourtzinis et al., 2017; Bheemanahalli et al., 2022a). The decrease in seed weight under combined stress resulted in a tradeoff between protein and oil content and was attributed to the reduced availability of substrate. Moreover, the study revealed that heat stress during grain development could lead to a reduction in lipid unsaturation levels, negatively impacting the nutritional quality of soybeans by reducing essential fatty acids such as linoleic acid and linolenic acid (Bukowski and Goslee, 2024). A decline in polyunsaturated fatty acid and linoleic acid under combined heat and drought stress accompanied an increase in oleic acid (Bellaloui et al., 2015). R01-416F and R15-2422 cultivars displayed the highest linoleic acid, contrasting with P46A86X and G4620RX, which exhibited the lowest. The cultivar with the least linoleic acid content had reduced yield and higher oleic acid (Figure 4). This alteration in linoleic and oleic acid was found to be attributed to the activation of the Triacylglycerol (TAG) degradation pathway, specifically the reduced activity of desaturase enzymes under combined stress conditions (Fehr, 2007; Bellaloui et al., 2015; Assefa et al., 2018; Kanai et al., 2019). While increased oleic acid levels and decreased linoleic and linolenic acid levels enhance the oxidative stability of the oil and result in more acceptable flavor quality scores, it’s important to note that these polyunsaturated fatty acids are known for their ability to lower cholesterol levels in human blood, thereby reducing the risk of heart diseases (Agyenim-Boateng et al., 2023). Consequently, a potential trade-off exists between improving oxidative stability and preserving soybean nutritional quality. To enhance soybean quality without compromising maturation and germination, ongoing research efforts are primarily directed toward manipulating lipase activity (Kanai et al., 2019). The cultivar with the highest yield (4775E3S) had higher seed protein under combined heat and drought than control but reduced oil content. This aligns with the previous studies, which show that drought and heat combination results in high seed protein content, providing an advantage to legumes over cereals in tolerating these stressors (Cohen et al., 2021a). However, cultivars with superior physiological traits, R15-2422, showed a higher percentage reduction in protein than the control. This finding indicates that interactive stress further increases the complexity of a trade-off in yield and quality traits (Figure 8).

### 4.5 Individual and combined treatments revealed unique traits and relationships

When plants experience simultaneous heat and drought stresses, their adaptation strategy is not just the sum of individual responses. Rather, it is influenced by the interaction of these stresses, perceived by plants as a new and unique state of stress (Mittler, 2006; Pandey et al., 2015). This results in different adaptation strategies under combined stress compared to individual stress. In our study, correlation analyses explained the unique and shared traits relationship between individual and combined treatments (Figure 7). We observed that the combined heat and drought stress has been shown to affect the physiological processes more severely than the individual stresses. The physiological parameters (stomatal conductance, transpiration, canopy temperature, photosynthesis) did not show significant correlations between control or single and combined stress (Figure 7). In contrast to hundred seed weight and pod number, all yield-related parameters showed weaker correlations between individual and combined stress treatments. This indicates greater plasticity in traits response to individual and combined stress. Under combined stress, some responses were shared with drought or heat stress. For instance, drought and combination reduced the pod number, unaffected by heat stress. Similarly, heat and combined stress increased the number of aborted, wrinkled, and small seeds, whereas drought stress did not reduce seed size.

This suggests that selecting stress tolerance based on individual stress performance may not accurately predict resilience under combined stress conditions. Our study, in line with similar observations in tobacco (Rizhsky et al., 2002) and Arabidopsis (Rizhsky et al., 2004), found that plant responses to combined stress can’t be directly compared to conclusions drawn from individual stress. This is likely because the combination of drought and heat stress triggers different, sometimes conflicting, signaling pathways than the individual stresses (Rizhsky et al., 2004). This can lead to synergistic effects, potentially reflecting a highly elevated stress level in the combined scenario beyond the combined impact of single stresses. Therefore, understanding the individual and compound effects of stressors is crucial for tolerance selection.

## 5. CONCLUSION

The individual heat or drought and combined stress-induced genetic variability in physiological, yield, and quality traits were explored. Findings demonstrated that the soybean cultivars were more susceptible to combined heat and drought than the individual stresses. Our results revealed a complex interplay between cultivar adaptation and the combined stress. Contrary to prior assumptions, resilience to individual stressors did not consistently perform similarly under combined stress across all measured traits. This suggests that cultivar selection for multi-stress environments requires a multifaceted approach, considering specific stress combinations and their intricate impact on plant physiology and performance. The observed phenotypic traits indicated synergistic effects of combined drought and heat stress, implying an elevated stress level beyond the additive effect of single stressors. This research highlights the importance of understanding cultivar-specific responses to combined stresses for developing stress-tolerant cultivars.

## ACKNOWLEDGEMENTS

We appreciate the Plant Stress Physiology Lab team for assisting with data collection. We thank David Brand for his technical support during the experiment. Please note that mentioning specific brand names or products in this publication is for informational purposes only and doesn’t mean we endorse or recommend them.

## DATA AVAILABILITY STATEMENT

The datasets used and/or analyzed during the current study are available from the corresponding author upon reasonable request.

