## Supplemental Material for "Negative Synergistic Effects of Drought and Heat During Flowering and Seed Setting in Soybean"

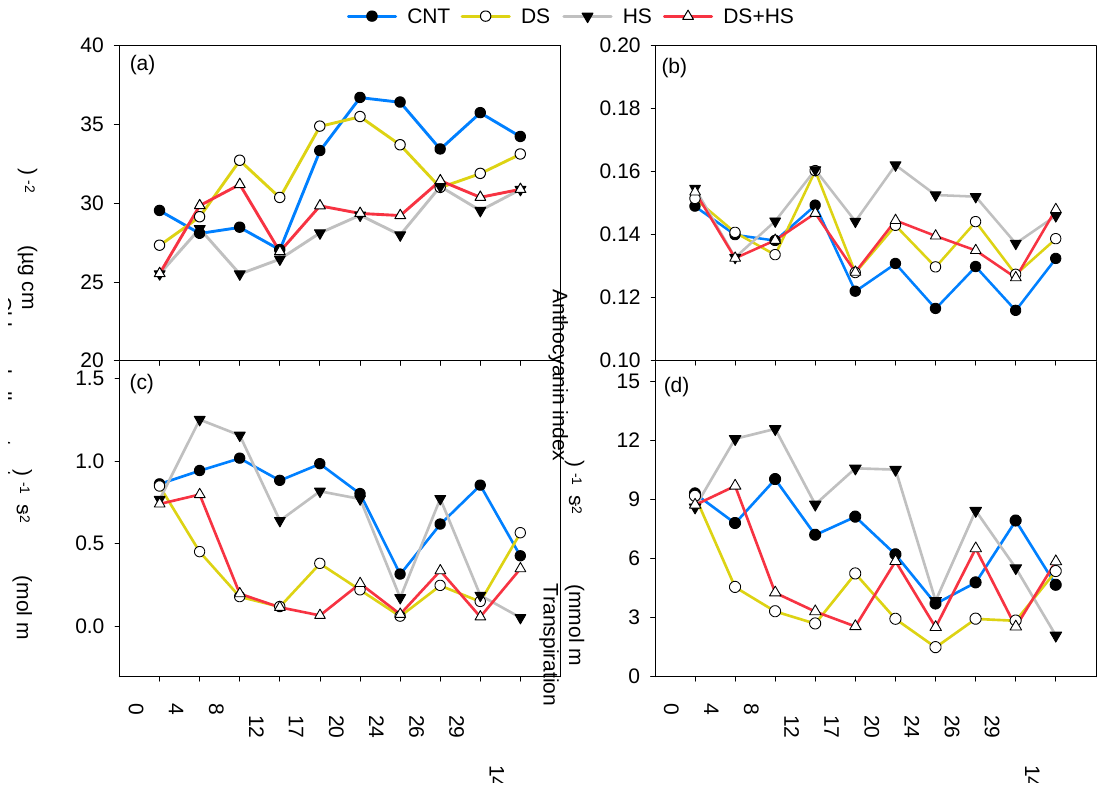
Supplementary Figure 1. Temporal distribution of chlorophyll content (µg cm^-2^, a), anthocyanin index (b) stomatal conductance (mol m^-2^ s^-1^, c), and transpiration (mmol m^-2^ s^-1^, d) under drought, heat, and their combination during the flowering seed setting stage of soybean cultivars Vertical bars denote SE. CNT - control, DS - drought stress, HS - heat stress, and DS+HS - combined heat and drought stresses. 14 DAR- 14 days after stress.

**Supplementary Table 1** List of ten soybean cultivars used in the experiment.

| Cultivar | Brand | Maturity group | Remark |
| --- | --- | --- | --- |
| 44-D49 | Armor | 4 | Roundup Ready 2 Xtend |
| 4775E3S | Progeny Ag | 4 | Enlist trait |
| DG4825RR2/STS | Delta Grow | 4 | Late Roundup Ready |
| DM 45X61 | Donmario | 4 | Roundup Ready 2 Xtend |
| G4620RX | AgriGold | 4 | Roundup Ready 2 Xtend |
| LS5009XS | Local Seed | 5 | Roundup Ready 2 Xtend |
| P46A86X | Pioneer | 4 | Roundup Ready 2 Xtend |
| S48XT90 | Dyna-Gro | 4 | Roundup Ready 2 Xtend |
| R01-416F | High protein | 4 | Improved nitrogen fixation under drought |
| R15-2422 | University of Arkansas | 4 | Advanced breeding line |

**Supplementary Table 2** List of the vegetation indices and their equations used in this study.

| Vegetation index | Equation | Reference |
| --- | --- | --- |
| Chlorophyll index of green (CIgreen) | $\frac{R_{NIR}-R_{G}}{R_{G}}$ | (Gitelson et al., 2003) |
| Chlorophyll index of red-edge (CIred-edge) | $\frac{R_{NIR}-R_{RE}}{R_{RE}}$ | (Steele et al., 2008) |
| Chlorophyll vegetation index (CVI) | $\frac{R_{NIR}*R_{R}}{R_{G}*R_{G}}$ | (Vincini et al., 2008) |
| Normalized difference red-edge index (NDRE) | $\frac{R_{NIR}-R_{RE}}{R_{NIR}+R_{RE}}$ | (Thompson et al., 2019) |
| Transformed Chlorophyll  Absorption In Reflectance Index (TCARI) | $3\times[\left( R_{700}-R_{670} \right)-0.2\times(R_{700}-R_{550})\times\frac{R_{700}}{R_{670}}$ | (Haboudane et al., 2002) |
| Visible Atmospherically  Resistant Index (VARI) | $\frac{R_{550}-R_{660}}{R_{550}+R_{660}-R_{470}}$ | (Gitelson et al., 2002) |

**Supplementary Table 3** Primers for Real-time quantitative PCR.

| **gene** | **primer ID** | **sequence(5'to3')** |
| --- | --- | --- |
| GLYMA.02G276600/ELF1B | Forward | GGTGATGAGACAGAGGAAGATAAG |
|  | Reverse | GCTTAACATCGAGAAGGACAGA |
| GLYMA.10G23600 | Forward | CCTACATCTGAGCAACTCGAATA |
|  | Reverse | GACGAGGTTCTTCTCCCATAAC |
| GLYMA.07G109100 | Forward | CCATCACTTCCCAAACAAACAC |
|  | Reverse | TTGGCGATGGAGTTGAAGAG |
| GLYMA.03G30040 | Forward | CTGGAGAGTCCATCAAGGAAAC |
|  | Reverse | CCGTCATTCTTTCTGCCTTCT |
